# Dual pathway architecture in songbirds enables robust sensorimotor learning

**DOI:** 10.64898/2026.05.07.723469

**Authors:** Remya Sankar, Atharv Suryawanshi, Nicolas P. Rougier, Arthur Leblois

**Author notes:** **For correspondence:** (AL); (NPR). These authors contributed equally to this work (Nicolas P. Rougier, Arthur Leblois).

## Abstract

The acquisition of sensorimotor skills critically depends on basal ganglia (BG)–thalamo–cortical circuits. Prevailing theories propose that the BG optimize motor output through reinforcement learning (RL), using internal performance evaluations to approximate stochastic gradient ascent. However, this framework struggles in non-convex performance landscapes, where local optima hinder efficient learning. Songbirds provide a compelling biological example of robust sensorimotor learning, mastering complex vocalizations through trial-and-error within a specialized BG–thalamo–cortical architecture. Here, we present a computational model constrained by the anatomy, physiology, and developmental trajectory of the zebra finch song system. The model combines a BG-driven RL pathway with a parallel cortical motor pathway that progressively consolidates successful motor patterns via Hebbian plasticity. In addition, we incorporate synaptic volatility within the BG pathway, introducing structured variability across learning. Through simulations of vocal learning using both a biophysical syrinx model and synthetic performance landscapes, we demonstrate that this dual-pathway architecture reliably converges to global optima and outperforms standard and noise-annealed RL approaches. The model reproduces key experimental features of song learning, including non-monotonic learning trajectories, a gradual reduction in motor variability, and the developmental transfer of motor control from subcortical to cortical circuits. Mechanistically, delayed maturation of the cortical pathway provides an implicit regulation of the exploration–exploitation trade-off, while synaptic volatility enables escape from local optima. These results highlight the importance of neural circuit architecture and dynamics in efficient learning, and suggest biologically inspired design principles for improving the robustness and sample efficiency of artificial RL systems in complex sensorimotor domains.

## Introduction

Acquisition and maintenance of sensorimotor skills are based on optimizing motor output based on internal evaluations of the behaviors performed and their results (***Hikosaka et al., 2002***). In the brain, this process is heavily dependent on basal ganglia (BG)–thalamo–cortical loops (***Hikosaka et al., 2002***; ***Graybiel and Grafton, 2015***). A prominent view frames BG–cortical interactions as a form of reinforcement learning (RL), in which dopamine influx in the BG signals performance (***Schultz et al., 1997***; ***Kasdin et al., 2025***) and dopamine-dependent cortico-striatal plasticity strengthens motor programs that previously yielded better-than-expected outcomes (***Wickens et al., 2003***; ***Doya, 2000***). Over many trials, this mechanism approximates stochastic gradient ascent on performance (***Williams, 1992***). However, when applied to continuous embodied sensorimotor tasks, the gradient-based RL framework faces major limitations. High-dimensional state–action spaces (***Bellman, 1957***), nonlinear body dynamics (***Bernstein, 1967***), and complex performance landscapes make gradient ascent slow, sample-inefficient, and vulnerable to local optima (***Peters and Schaal, 2008***; ***Dhawale et al., 2017***). Artificial RL agents typically rely on heuristic exploration strategies (e.g. *ϵ*-greedy or uncertainty bonuses), (***Sutton and Barto, 2018***; ***George and Powell, 2006***) that lack biological grounding and do not resemble the richer exploratory behaviors observed in humans or animals (***Gershman and Daw, 2017***; ***Shen and Dayan, 2025***).

Juvenile songbirds provide a model system that exposes these shortcomings. Learning to imitate a tutor’s song requires exploring a vast sensorimotor space to approach an optimal performance (***Mooney, 2009***; ***Fee and Goldberg, 2011***). Yet, the trajectory of learning is highly irregular and non-monotonic. Performance often deteriorates after sleep before improving with practice, and the magnitude of this post-sleep deterioration predicts later imitation quality (***Derégnaucourt et al., 2005***). Moreover, changes within the day, overnight fluctuations, and slower consolidation on a week scale display a partial misalignment (***Kollmorgen et al., 2020***), making song learning an erratic process that defies simple gradient-ascent interpretations. These saltatory transitions suggest that the brain employs mechanisms beyond straightforward local optimization, allowing discontinuous traversals across rugged performance landscapes.

Songbirds rely on a dual pathway brain circuitry dedicated to vocal learning for song acquisition and maintenance. A cortical-like motor pathway and a BG-thalamo-cortical circuit interact during learning, with structured variability, pathway-specific plasticity, and delayed maturation providing a substrate for guided exploration and consolidation (***Mooney, 2009***). The motor pathway comprising HVC (used as a proper name) and the robust nucleus of the arcopallium (RA) executes vocal output, generating the precise timing, sequencing, and motor patterns driving the song (***Hahnloser et al., 2002***; ***Sober and Brainard, 2009***). During development, RA firing evolves from variable juvenile patterns to highly stereotyped adult burst sequences aligned to the song with 10-20 ms precision (***Ölveczky and Gardner, 2011***). In parallel, a BG–thalamo–cortical loop (Area X → DLM → LMAN → Area X) receives dense dopaminergic input, evaluates performance (***Kasdin et al., 2025***), and injects variability and corrective bias into RA through LMAN (***Warren et al., 2011***; ***Andalman and Fee, 2009***; ***Kao et al., 2005***). This BG-derived bias is gradually consolidated into the increasingly influential motor pathway through activity-dependent plasticity (***Aronov et al., 2008***; ***Mehaffey and Doupe, 2015***). Toward the beginning of learning, the BG pathway dominates exploration, while direct cortical motor projections strengthen over time, eventually making BG input unnecessary to produce the crystallized song (***Bottjer et al., 1984***; ***Scharff and Nottebohm, 1991***), though its removal reduces residual variability (***Dhawale et al., 2017***; ***Kao et al., 2005***).

To explore how biological systems solve complex sensorimotor problems, we develop a computational model that captures vocal learning in a way consistent with and constrained by the behavioral, physiological, and anatomical characteristics of song learning in zebra finches (Figure 1a,b). In this model, we propose a neural mechanism inspired by the song system that addresses key limitations of standard RL. Through simulations of the neuronal network driving vocal learning across diverse performance landscapes, we show that the interaction between circuit architecture and synaptic plasticity enables the system to escape local optima and more consistently reach global solutions compared to standard RL algorithms. We further propose that synaptic volatility and delayed maturation of the cortical pathway could produce the observed non-monotonic learning features such as post-sleep deterioration while ensuring optimal final performance. Altogether, this work highlights architectural principles that could enhance the robustness and sample efficiency of RL for real-world sensorimotor control.

**Figure 1.**
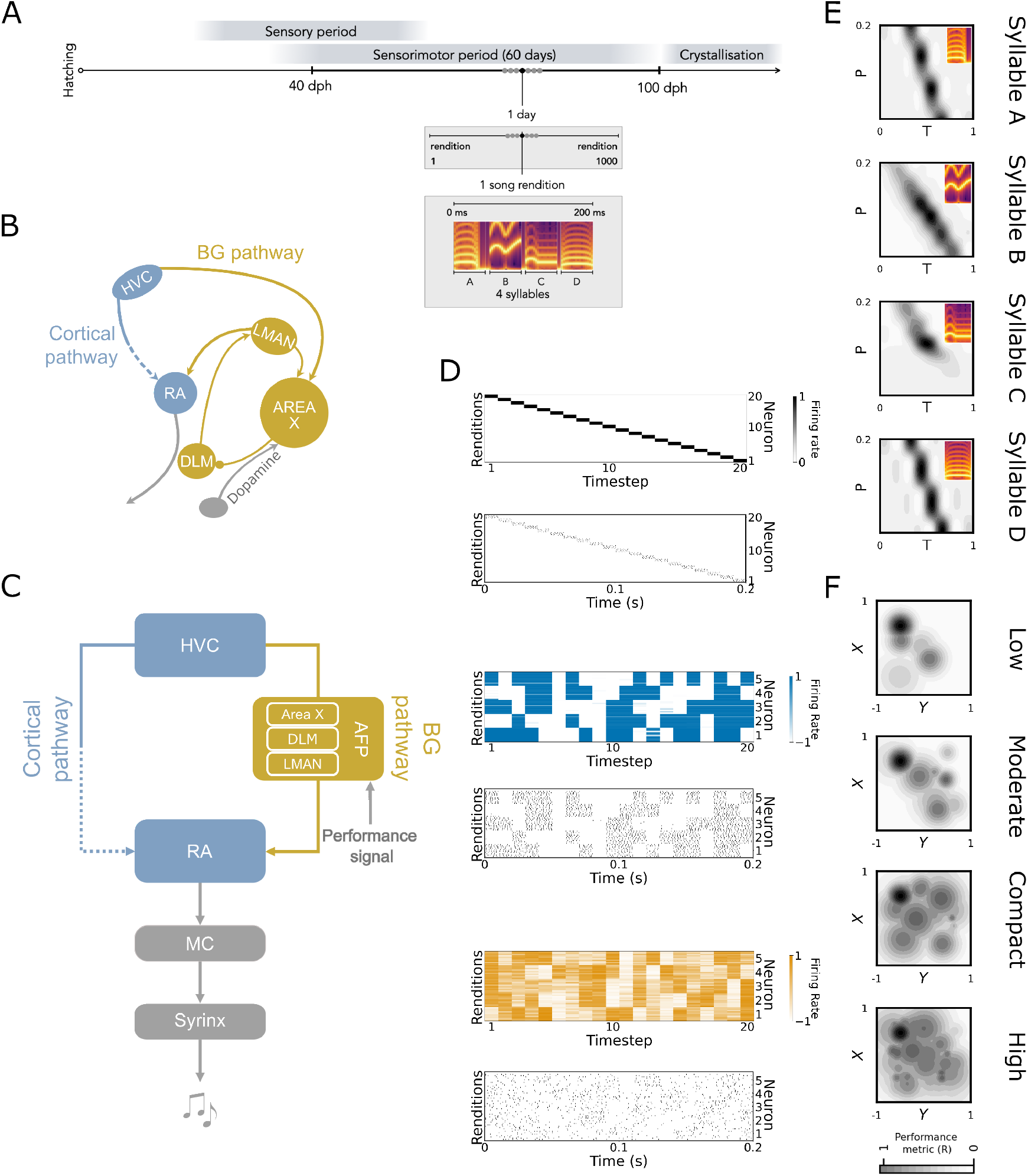
The dual pathway model for vocal learning. **A**. Timeline of song learning. The model learns to produce 4 syllables (50 ms each) per motif, and practices the motif 1000 times per day for 60 days (dph: days post hatch), similar to the sensorimotor phase of zebra finches. **B**. The neural substrates involved in vocal learning in zebra finches, the song system. The cortical pathway driving adult song production, in blue, with dotted lines representing its delayed maturation and gradual strengthening over the sensorimotor phase. The subcortical pathway, in yellow, necessary for song learning and maintenance, receiving information about performance quality through dopaminergic input. **C**. The architecture of the dual pathway model is derived from the song system. The cortical pathway, in blue, consists of two nuclei, HVC (used as a proper name) and the robust nucleus of the arcopallium (RA). The pathway strengthens during learning (dotted). The subcortical pathway, in yellow, consists of one basal ganglia (BG) neuronal population, representing the BG-thalamo-cortical loop (comprising Area X, medial part of the dorsolateral nucleus (DLM) and lateral magnocellular nucleus of the anterior neostriatum (LMAN)). The motor control (MC) layer transforms the RA neural commands to the appropriate vocalisation parameters (see Methods. **D**. Representative neural activity in each neuronal population of the model at the end of the sensorimotor phase. For each population, the top panel shows the firing rate generated for each 10 ms time bin during song, and the bottom panel shows illustrative spiking activity (see Methods), where each vertical black line denotes a spike. HVC neurons (black) fire sequentially, providing a timing scaffold for the song. By the end of the sensorimotor period, RA neuronal populations (blue) display sparse, stereotyped bursts across renditions. BG neuronal populations (yellow) exhibit high rendition-to-rendition variability in firing rate. **E**. Examples of performance landscapes generated using a biophysical model of the syrinx (see Methods) with four target syllables (inset). The performance metric at each point (determined by pressure, P, on y-axis, tension, T, on x-axis) on the landscape is indicated by the grey heatmap. The landscapes have a few global optima (performance metric ≈ 1) and several local optima (performance metric < .7). **F**. Examples of Gaussian-based landscapes from each density category. The landscape is generated by superimposing several Gaussian functions (see Methods). The superimposition results in several local (performance metric < .7) and one globally optimal solution (performance metric = 1). The landscapes are divided into four classes according to the number of local optima present: low (2-5), moderate (3-9), compact (8-18), high (11-25).

## Results

### The dual pathway model

We investigate the neural mechanisms of song learning using a computational model of the song-related network constrained by the anatomy and physiology of the song system described in the Introduction (Figure 1a-c). The model includes three populations, each representing the contribution of a song system nucleus or pathway to song acquisition: one population representing nucleus HVC, one representing nucleus RA, and a BG population representing the contribution of the basal ganglia-thalamo-cortical circuit. HVC and RA are connected via two pathways: a direct cortical (HVC-RA) pathway and a polysynaptic pathway through BG.

Song learning is simulated over a 60-day sensorimotor period mimicking the sensorimotor period of song learning, with 1,000 motif renditions per day (Figure 1a, ***Tchernichovski et al. (2000***)). HVC provides a fixed, time-locked representation of the song motif sequence (Figure 1d, ***Hahnloser et al. (2002***)), assumed for simplicity to be already in place by the beginning of the sensorimotor learning (***Mackevicius et al., 2023***). Initially, HVC–RA synapses are absent and gradually strengthen during learning (***Mooney and Rao, 1994***), while HVC–BG connectivity is random. RA activity drives a 2D motor output controlling vocal production, which is evaluated at each time step (see Methods) to generate a dopaminergic performance prediction error signal (***Gadagkar et al., 2016***; ***Kasdin et al., 2025***). Performance is assessed either through a biophysical syrinx model, comparing produced vocalizations to tutor syllables (Eq 17, see Methods, Figure S1a, Figure 1e, ***Titze and Martin (1998***); ***Amador and Margoliash (2013***)), or via synthetic landscapes with controlled numbers of local optima (Figure S1c, d, Figure 1f). In both cases, a performance prediction error signal calculated from current and past performance as Eq 10 is fed into a tri-partite reinforcement learning plasticity rule at HVC-BG synapses following Eq 9. This plasticity rule is known to drive gradient ascent in the performance landscape (***Williams, 1992***). It will allow BG output, and thereby RA output, to gradually approach the closest local maximum in the performance landscape. This gradient ascent strategy is clearly insufficient and bound to fail in landscapes with multiple sub-optimal maxima such as the ones derived from the syrinx model (Figure 1e) or designed artificially to test our model (Figure 1f).

To overcome the limitations of gradient-based learning in multi-peaked landscapes, the model incorporates two key biological features. First, Hebbian plasticity in the HVC–RA pathway progressively consolidates BG-driven motor patterns (Eq 12, ***Mehaffey and Doupe (2015***); ***Teşileanu et al. (2017***)). Second, synaptic volatility in HVC–BG connections introduces overnight perturbations (Eq 13), mimicking the continuous remodeling of the cortico-striatal synapses with volatility over hours to days (***Kuo and Liu, 2019***; ***Cepeda et al., 2003***; ***Holtmaat and Svoboda, 2009***). This volatility induces post-sleep deterioration (***Derégnaucourt et al., 2005***), promoting non-monotonic learning and exploration. Together, these mechanisms enable learning in non-convex performance landscapes. In the following sections, we compare the model’s behavior to songbird data, analyze the interaction between pathways, and benchmark its performance against alternative learning strategies.

### Song imitation with an artificial syrinx or in arbitrary performance landscapes

In Figure 2, we evaluate the dual-pathway model on a motor learning task involving vocal production through a biophysical syrinx model (see Figure 1). The network learns to generate RA activity patterns that reproduce four tutor syllables (each 50 ms duration) forming a song motif (Figure 2a) by maximizing the similarity between produced and target vocalizations. Initially, outputs lie far from optimal regions of the performance landscape, but through exploration the model converges to accurate imitations (Figure 1b-e). This improvement is accompanied by systematic exploration of motor parameters controlling syrinx dynamics. Across 100 simulations, individual syllables were successfully imitated (terminal performance > 0.7) in over 85% of cases, with high median performance (median terminal performance^1^ of A: 0.93 [0.92-0.97], B: 0.98 [0.95-0.99], C: 0.98 [0.98-0.99], D: 0.98 [0.97-0.99]) (Figure 2f), and full motifs were successfully learned in more than 95% of simulations. Failures (≈5%) corresponded to convergence to suboptimal local maxima.

**Figure 2.**
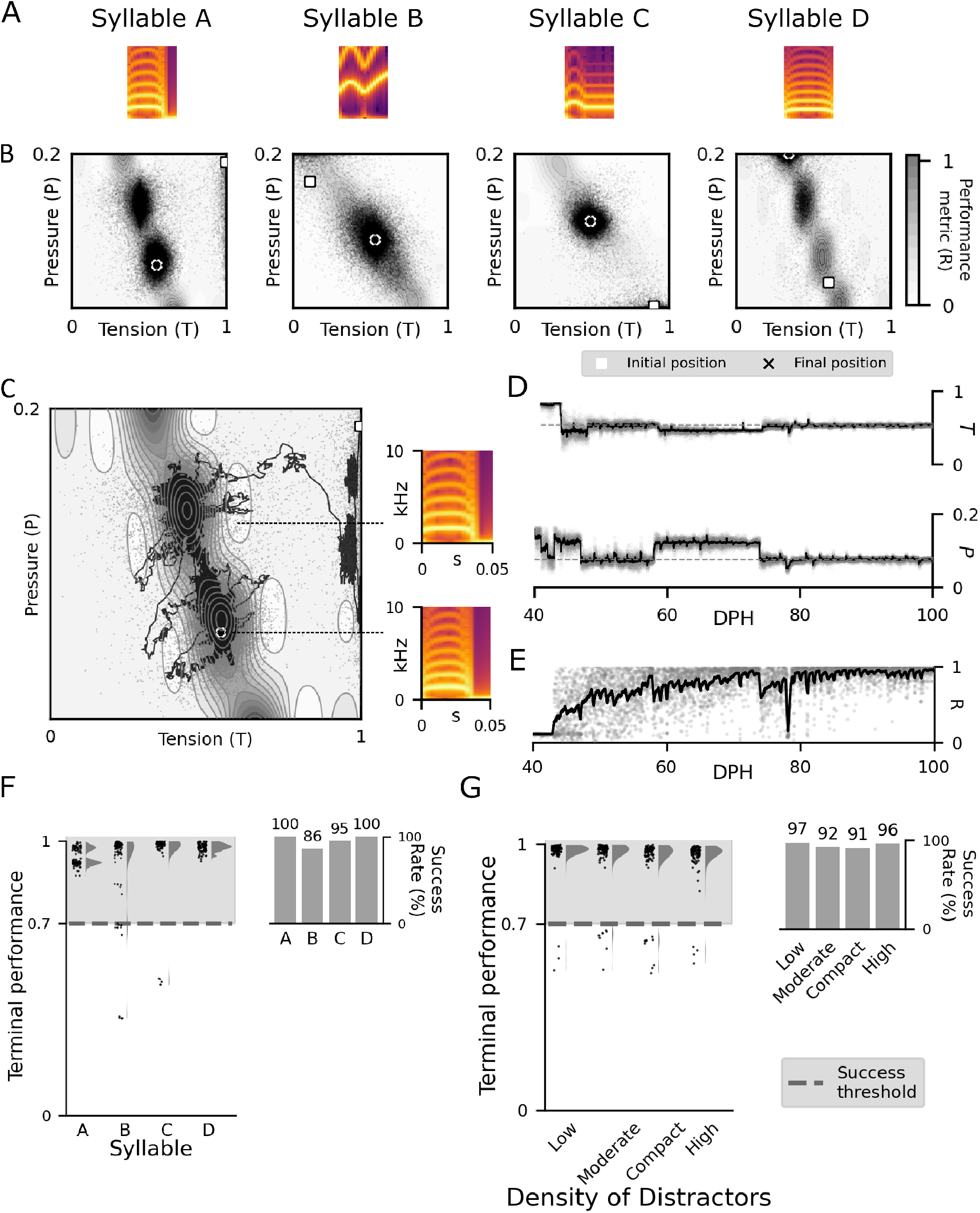
Song acquisition in the dual pathway model. **A**. Target song composed of 4 syllables. **B**. Model simulation of the learning process for each syllable over the sensorimotor period. The model starts at a random initial position (square) and is able to find the optimal solution (cross) over the sensorimotor period of 60 days with 1000 iterations each day. The output at each iteration is shown as a black dot. Each panel shows the motor outputs produced for the corresponding syllable. **C**. Example trajectory of the model output. The two-dimensional MC output is fed as pressure and tension in the syrinx model, driving the production of each syllable. The model starts at a random initial position (square), produces several sub-optimal vocalisations, over the coarse of learning (smoothened trajectory in black) and eventually finds the globally optimal solution (cross). Two example output vocalisations are shown on the right, corresponding to a locally (top) and globally (bottom) optimal performance, with differing fundamental frequencies. **D**. Smoothened trajectories for the control parameters representing the syrinx pressure and tension for the example syllable simulation in C are shown in black. The target output for each control parameter is shown in grey horizontal dotted lines. The model explores different outputs (grey dots) in the two dimensions in the beginning of sensorimotor learning and eventually converges at the target output. **E**. Smoothened trajectory for the performance metric obtained by the model at every rendition of the example syllable over the sensorimotor period (in black). Grey dots represent performance metric at each rendition. **F**. Performance of the model over 100 simulations on the syrinx-based landscapes. Left: Terminal performance for all simulations (black dots), calculated as the average performance over the last 100 trials on the last day (see Methods). Performance metrics above the threshold (shaded portion above the black dotted line), lie close to the global optimum. The violin plot denotes the distribution of the terminal performance (black dots) across the 100 simulations. Right: Success rate for each syllable, i.e. the proportion of simulations that learned the optimal performance (achieved terminal performance > .7). **G**. Performance of the model over 100 simulations on the gaussian-based landscapes, with increasing density of local optima. Same conventions as in F.

We then assessed performance on synthetic landscapes with a single global optimum and varying densities of local optima (Figure 1f). As distractor density increased, success rates declined modestly and plateaued, reflecting increased trapping in local optima (Figure 2g). At high densities, performance improved again, as closer spacing between optima facilitated transitions toward the global solution. Overall, these results demonstrate that the dual-pathway model achieves robust learning across a wide range of landscape complexities.

### Subcortical circuitry is required for vocal learning by the dual pathway model

To disentangle the roles of the cortical (HVC–RA) and subcortical (HVC–BG–RA) pathways across learning, we performed in silico lesions mimicking experimental manipulations in songbirds. Consistent with prior studies, we selectively inactivated BG inputs to RA at different developmental stages to assess their contribution to learning and production. Early BG inactivation (before or at 40 dph) resulted in low imitation quality and limited improvement over time (Figure 3a, left panel), with reduced motor variability, indicating that the cortical pathway alone cannot support learning at this stage. Mid-stage inactivation (70 dph) led to moderate performance impairments (success rate drops from 91% to 80%) and a sharp drop in variability (from 0.11 [0.09-0.14] to 0.03 [0.02-0.04]), with learning plateauing thereafter (Figure 3 a,b,c, middle panels). In contrast, late inactivation (100 dph) had minimal impact on vocal output and performance (0.98 [0.97-0.99] vs 0.98 [0.96-0.99], Figure 3a-c), with only a slight reduction in variability (from 0.04 [0.03-0.06] to 0.01 [0.002-0.01]), showing that BG input becomes largely dispensable once learning has converged in the model, as shown experimentally (***Bottjer et al., 1984***; ***Scharff and Nottebohm, 1991***; ***Aronov et al., 2008, 2011***).

**Figure 3.**
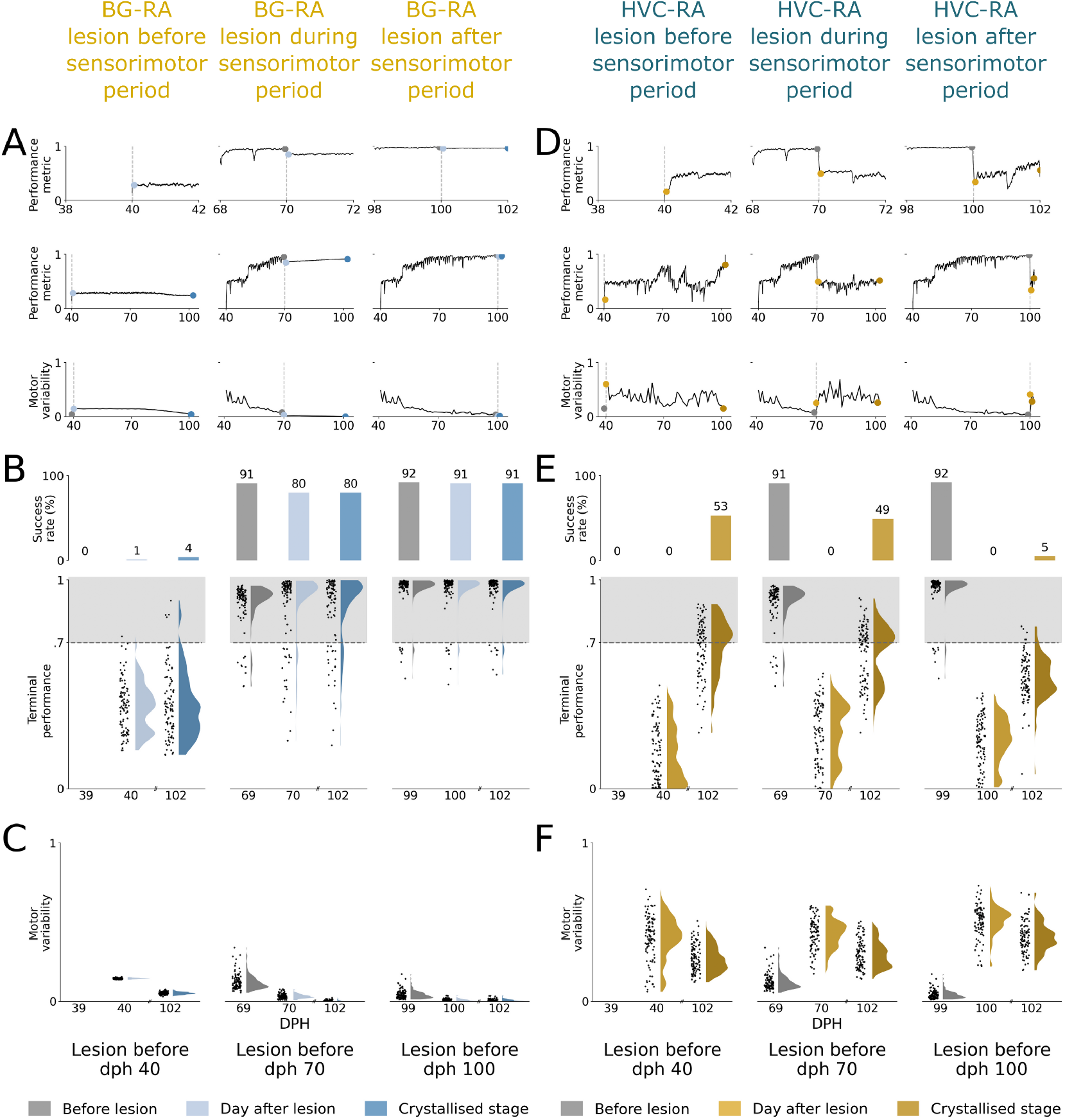
Effects of inactivating the BG and cortical inputs to RA, respectively, at different stages of the sensorimotor learning period. **A**. Performance before and after inactivation of BG input to RA at different stages of sensorimotor learning in representative simulations. The three columns show the effect of BG inputs being inactivated before (left), during (middle) and after (right) sensorimotor learning. Top panel: Performance (black) two days before and after lesion. Vertical dotted line denotes the lesion time. Middle panel: Performance (black) throughout the sensorimotor period. Bottom panel: Motor variability over the sensorimotor period. In all three panels, the grey dots denote the performance metric value at the end of the day before inactivation. The light blue dots denote the metric at the start of day after inactivation. The blue/yellow dots denote the metric at the end of the sensorimotor period. **B**. Distribution of success rate (top) and terminal performance (bottom) across 100 simulations before and after inactivation of BG input to RA performed at different stages of sensorimotor learning. Bottom: Terminal performance for all simulations (black dots, see Methods). Performance metrics above the threshold (shaded portion above the black dotted line), lie close to the global optimum. The violin plot denotes the distribution of the terminal performance (black dots) across the 100 simulations. Top: Success rate for each condition (proportion of simulation achieving terminal performance > 0.7). **C**. Motor variability across 100 simulations before and after inactivation of BG input to RA performed at different stages of sensorimotor learning. Black dots show the motor variability for each simulation (see Methods) while violin plot denotes the distribution of motor variability across the 100 simulations. **D-F**. Corresponds to A-C for the suppression of HVC input into RA (in yellow).

Conversely, inactivating the cortical pathway (HVC–RA) at any stage prevented stable performance despite ongoing exploration (Figure 3d-e). At mid and late stages, this caused a dramatic drop in performance (dph 70: from 0.92 [0.89 - 0.94] to 0.28 [0.14 - 0.39] and dph 100: from 0.98 [0.96 - 0.99] to 0.24 [0.15 - 0.31], success rate falling to 0%), and a strong increase in variability (dph 70: from 0.11 [0.09 - 0.15] to 0.44 [0.36 - 0.49] and dph 100: from 0.04 [0.03 - 0.06] to 0.53 [0.46 -0.57]), reflecting unopposed BG-driven fluctuations. The BG pathway is thus essential for early exploration and learning but becomes unnecessary for stable production after crystallization, while the cortical pathway progressively assumes control and suppresses variability. BG lesions also abolish overnight performance deterioration in the model, as it is driven by BG-mediated synaptic volatility. Together, these results support a dual-pathway mechanism in which BG-driven exploration enables learning, while cortical circuits consolidate stable motor patterns.

### Neural activity patterns in the model parallels song system neurophysiology

The model reproduces key neurophysiological features of the song system. HVC exhibits sparse, time-locked burst firing, with each time point corresponding to a distinct target output and performance landscape (Figure 4a). HVC neurons project to both RA and the basal ganglia (BG), forming parallel pathways. Throughout learning, BG activity remains variable across renditions (dph 40: 0.19±0.09; dph 99: 0.17±0.05; mean±std, Figure 4c), while RA dynamics evolve markedly (Figure 4b). Early in learning, weak HVC–RA synapses make RA activity largely driven by the BG pathway, resulting in high variability (dph 40: 0.36±0.29; mean±std). As HVC–RA synapses strengthen, RA becomes increasingly driven by stable HVC input, reducing sensitivity to BG fluctuations and leading to lower variability (dph 100: 0.05±0.13; mean±std). This transition from variable to stereotyped RA activity mirrors experimental observations in juvenile zebra finches (***Ölveczky et al., 2011***) and reflects the progressive stabilization of motor output. Overall, the model captures both the structured temporal drive from HVC and the gradual refinement of activity underlying vocal learning, with behavior shifting from exploratory to stereotyped output.

**Figure 4.**
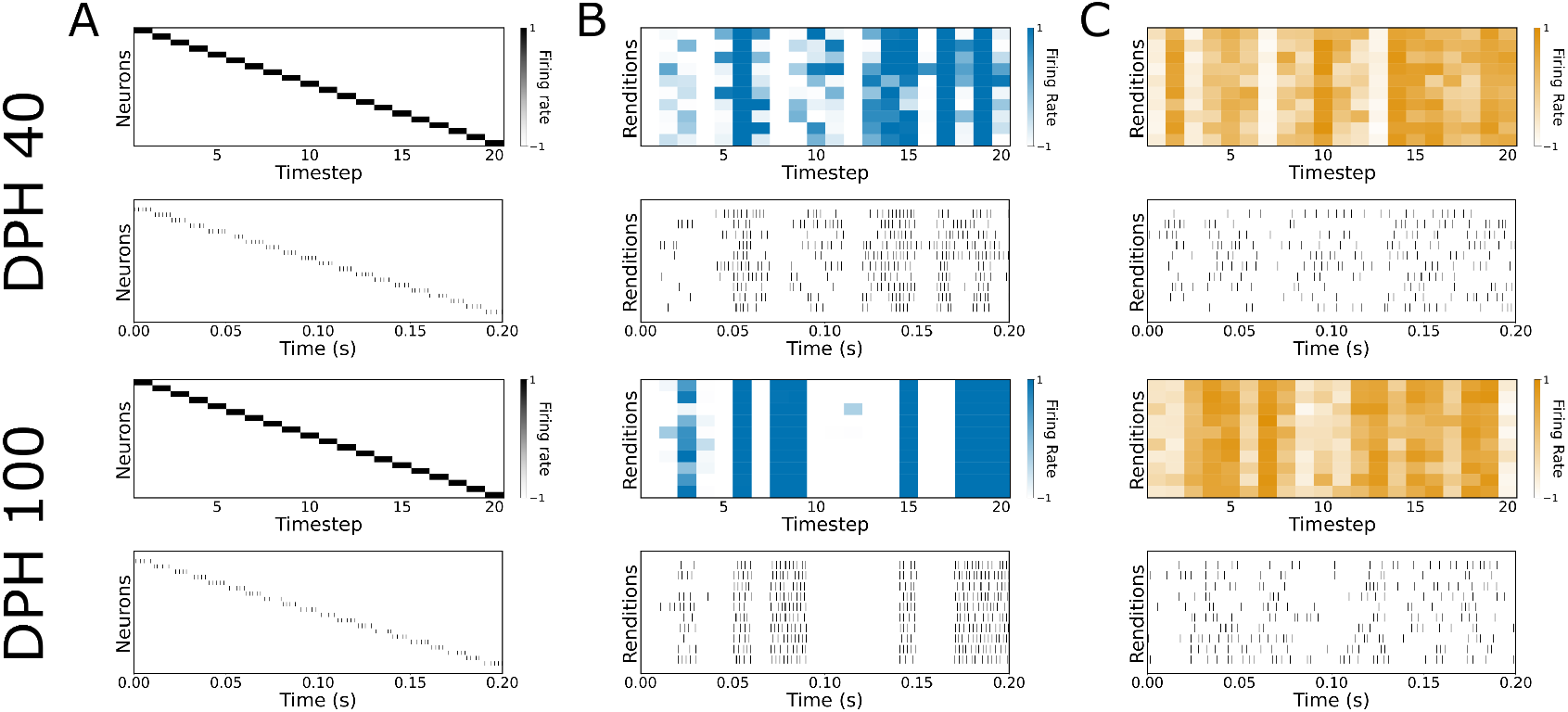
Neural activity patterns across the sensorimotor learning period (40 dph vs 100 dph) in the model. **A**. Neurons in the HVC layer encode time during song and provide context information to the model. The activity pattern of HVC neurons (black) across one song rendition. Top panel: Firing rate of multiple neurons across time steps during one song rendition. Bottom panel: Poisson spike trains generated based on the above firing rate profiles (see Methods). Each vertical black line corresponds to one action potential. **B**. Neurons in the RA layer have variable firing rates across song renditions in the beginning of sensorimotor learning and stereotyped patterns towards the end of sensorimotor learning. Top panel: Firing rate of one RA neuron (shades of blue) across ten song rendition. Bottom panel: Poisson spike trains generated based on the above firing rate profiles. **C**. The neurons in the BG layer have variable firing rates across song renditions throughout sensorimotor learning, akin to LMAN neural activity. Top panel: Firing rate of one BG neuron (shades of blue) across 10 song renditions. Bottom panel: Poisson spike trains generated based on the above firing rate profiles.

### Overnight performance deterioration aids sensorimotor learning

Juvenile zebra finches exhibit non-monotonic learning, with performance often deteriorating after sleep (***Derégnaucourt et al., 2005***). In our model, we introduce random synaptic perturbations at HVC–BG synapses during the night. The magnitude of this volatility is stochastic and modulated inversely by the extent of synaptic change accumulated during the preceding day (see section Meth- ods). It produces daily shifts in motor output. Without this volatility, the model follows local gradient ascent, converging monotonically to nearby suboptimal solutions and becoming trapped (Figure 5a–c). In contrast, with overnight volatility, synaptic changes are minimal during the day but substantially larger across nights (0.11 [0.05-0.21] vs 0.27 [0.09-0.51], Mann-Whitney U = 4.38e7, p < 0.001, r = 0.38), leading to systematic displacements in BG and RA activity. This induces broader, cross-day exploration of the performance landscape (Figure 5d–f), enabling the model to escape local optima and discover the global solution.

**Figure 5.**
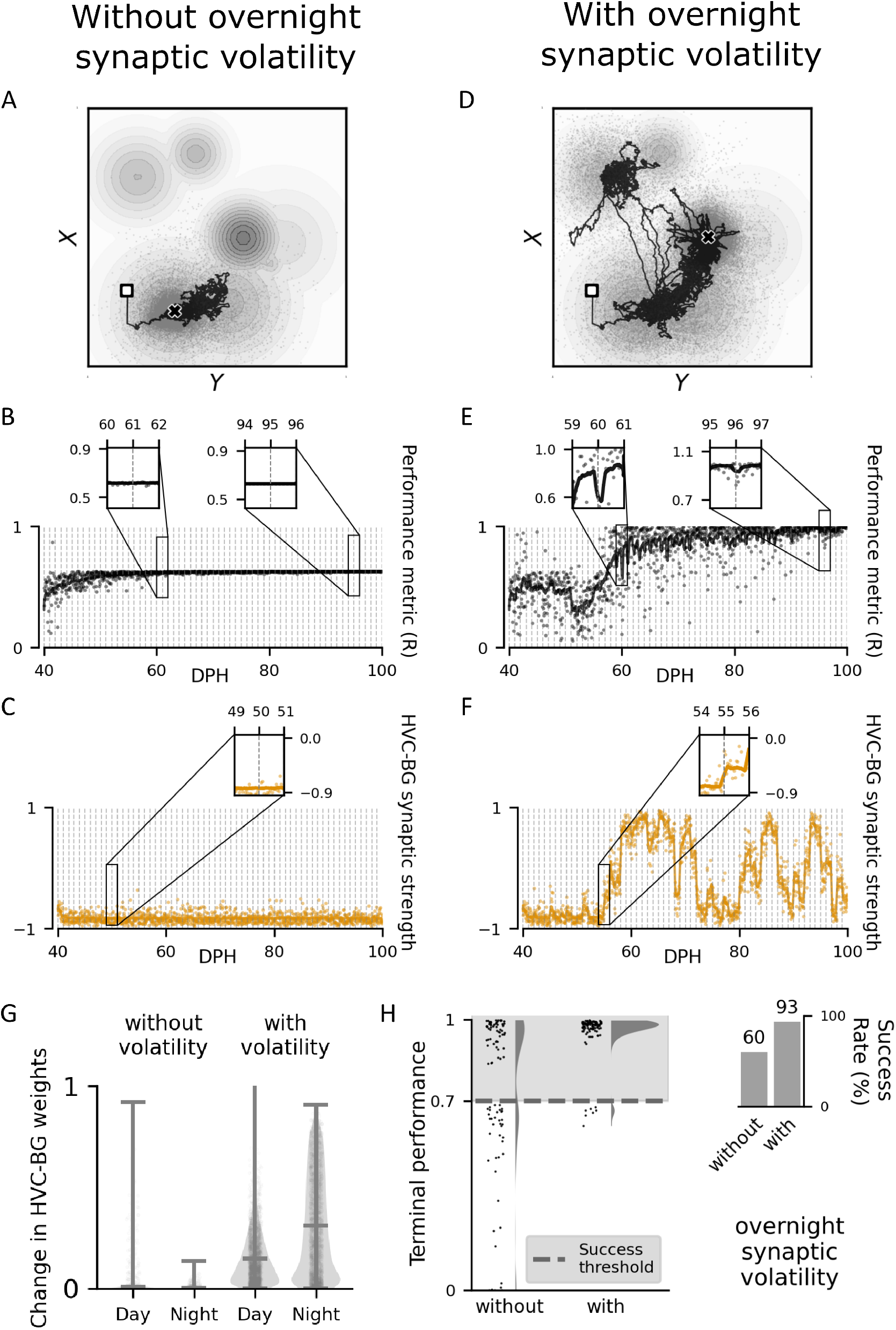
Synaptic volatility induces overnight displacement in performance and aids sensorimotor learning. The two columns show one representative example simulation of the learning of a single vocal gesture (single time step), with (right) and without (left) overnight synaptic volatility. **A**. Representative learning trajectory in the absence of synaptic volatility. Each black dot corresponds to one rendition of the motor output. The model converges (cross) at the sub-optimal solution closest to the initial position (square). **B**. Smoothened trajectory (black line) of the performance metric (black dots) over a representative simulation. Vertical grey lines denote day change. Inset: Performance metric over two days during the mid and late stages of sensorimotor learning displays a monotonic performance quality. **C**. Smoothened trajectory (yellow line) of the synaptic strength (yellow dots) of one representative HVC-BG weight. Vertical grey lines denote day change. Inset: Synaptic strength over two days shows gradual changes in the HVC-BG weight during the day and absence of overnight fluctuation. **D**. Representative learning trajectory with synaptic volatility, with same convention as in A. The model escapes sub-optimal solutions and converges (cross) at the globally optimal performance. **E**. Smoothened trajectory (black line) of the performance metric (black dots) over a representative simulation. Vertical grey lines denote day change. Inset: Performance metric over two days during the mid and late stages of sensorimotor learning displays gradual changes in performance during the day and sudden overnight fluctuations. The magnitude of overnight performance fluctuation reduces over the sensorimotor period. **F**. Smoothened trajectory (yellow line) of the synaptic strength (yellow dots) of one representative HVC-BG weight. Vertical grey lines denote day change. Inset: Synaptic strength over two days shows gradual changes in the HVC-BG weight during the day and strong fluctuations overnight. **G**. Change in HVC-BG synaptic strengths across the day vs across the night over one simulation with and and one without overnight synaptic volatility in the BG pathway. **H**. Terminal performance (left, black dots) and success rate (right) of the model over 100 simulations with and without overnight synaptic volatility in the BG pathway.

As a result, learning becomes non-monotonic across days: performance drops after each night but improves within each day, closely matching experimental observations (Figure 5e). This effect is strongest early in learning and diminishes over time. Across 100 simulations, success rates increased markedly with synaptic volatility (93% vs 60%, Mann-Whitney U = 1.07e9, p < 0.001, r = 0.408), along with higher terminal performance (0.98 [0.96-0.99] vs 0.90 [0.63-0.97], Figure 5h). These findings show that overnight synaptic volatility provides an effective exploration mechanism, facilitating escape from local optima and enabling robust learning in complex, non-convex landscapes.

### Delayed maturation of cortical pathway modulates exploration-exploitation transition

The dual-pathway model explores multiple regions of the sensorimotor landscape before converging to a stable solution (Figure 6b-d). Early in learning, the cortical pathway is weak and contributes little to motor output, which is instead dominated by the basal ganglia (BG) pathway. As a result, vocal output is highly variable and reflects BG-driven exploration. Over time, Hebbian plasticity strengthens the cortical pathway, progressively consolidating BG-led vocalizations. Cortical activity increasingly drives motor output, while the influence of the BG pathway diminishes. Consequently, RA activity and vocal output become more stable and less variable across development (Figure 6f).

**Figure 6.**
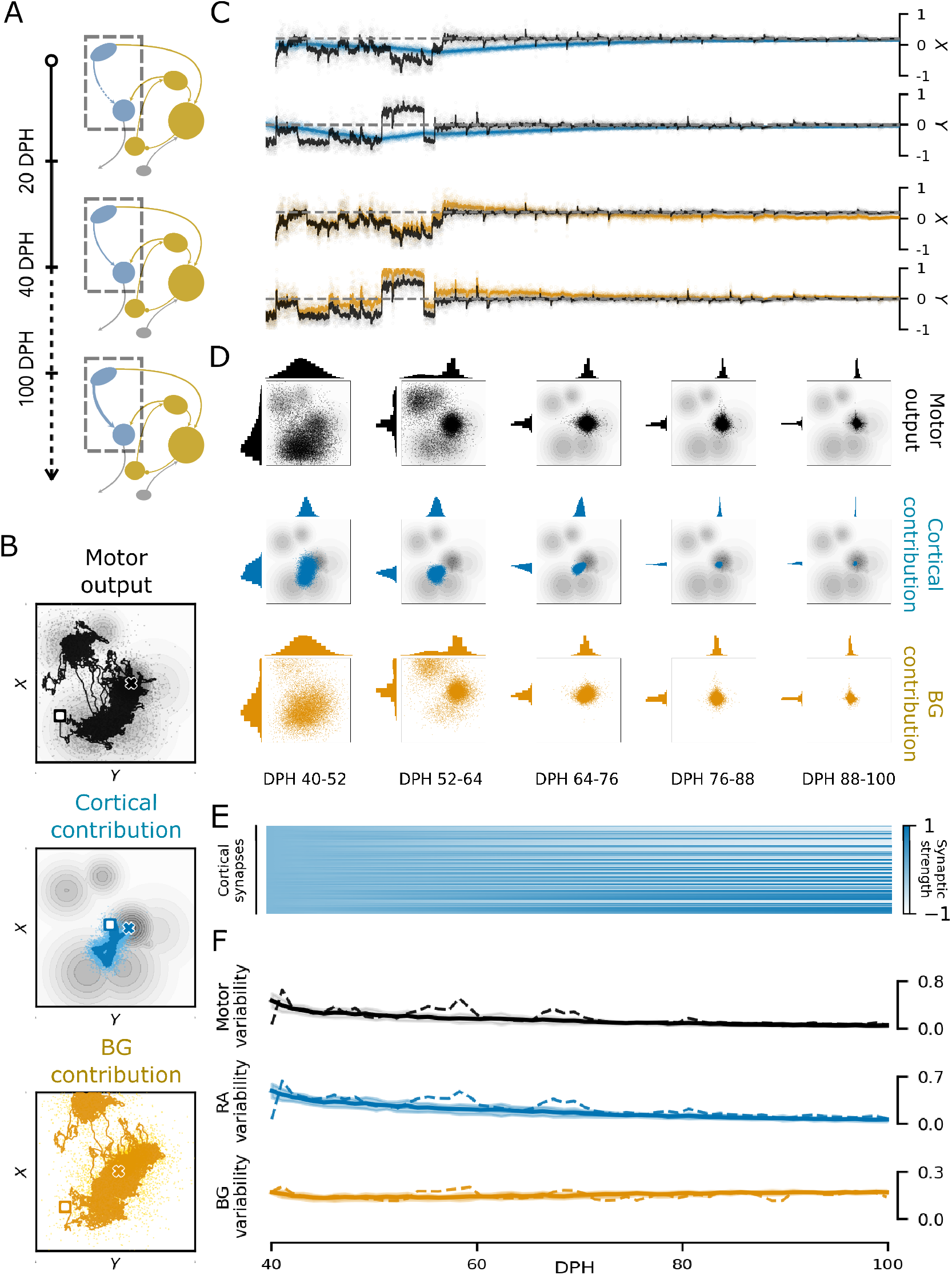
Delayed maturation of cortical pathway modulates the exploration-exploitation trade off. **A**. Schema of delayed maturation of the HVC projections to RA and its growing strength over development in male zebra finches. **B**. Representative trajectory of the model output for one time-step (single vocal gesture) in the dual pathway model. Smoothened trajectories of the overall motor output (top, black line), contribution of the cortical pathway to motor output (middle, blue line) and contribution of the BG pathway to motor output (bottom, yellow line). The cortical contribution slowly consolidates a trace of the BG-led exploration of the motor space. Squares denote initial positions and crosses denote final positions. **C**. Learning trajectory along each dimension. The overall motor output (black) overlaps primarily with the BG contribution (yellow) in the beginning of the sensorimotor phase. Over time, it aligns increasingly with the cortical pathway contribution (blue), as the model approaches convergence. As the motor output reaches the target output (grey dotted line), the BG contribution slowly fades to zero. **D**. Change in motor output across different stages of sensorimotor tasks (Top: overall output, middle: cortical pathway contribution, bottom: BG pathway contribution). The histogram show the distribution of motor output (dots) on the landscape in the two dimensions. BG contribution (bottom panel) initially induces a strong displacement in the overall motor output (top panel) as the cortical contribution (middle panel) incorporates a slow trace. However, its effect on motor output fades over time, eventually contributing to moderate exploration around the mainly cortical-driven output. **E**. Growth of synaptic strength (shades of blue) in the cortical pathway over the sensorimotor period (each line represents a single synapse). The HVC-RA synapses, initalized at zero weight, drive a strong net excitatory or net inhibitory effect on the postsynaptic RA populations at the end of learning. **F**. Top panel: Motor variability reduces over the sensorimotor period. Middle panel: Variability of RA activity reduces over the sensorimotor period. Bottom panel: Variability of BG activity remains stable over the sensorimotor period.

Importantly, this reduction in motor variability (dph 40: 0.41±0.12, dph 100: 0.04±0.02; mean±std) does not stem from changes within the BG itself, which remains consistently variable (dph 40: 0.17±0.01, dph 100: 0.17±0.01; mean±std). Rather, the impact of BG-driven fluctuations is atten-uated as the cortical pathway strengthens (Figure 6a,e), a process also reflected in the decreasing magnitude of overnight performance shifts (early: 0.4±0.29, late: 0.05±0.03; mean±std; Figure 5e). Overall, the model exhibits a gradual transition from BG-driven exploration to cortical-driven exploitation, reflecting a transfer of motor control from subcortical to cortical circuits. This shift emerges naturally from the delayed maturation of the cortical pathway, which implicitly regulates the exploration–exploitation trade-off and supports efficient sensorimotor learning.

### Dual pathway model surpasses alternate strategies for sensorimotor learning

We next assessed whether a dual-pathway architecture is necessary for robust learning by comparing it with single-pathway reinforcement learning (RL) models under different strategies. A standard single-pathway RL model (StdRL, see Section Methods) performed poorly on artificial landscapes, succeeding in fewer than one-third of simulations at low noise levels and remaining trapped in suboptimal solutions (Figure 7a). Increasing noise improved exploration and raised success rates, but excessive noise degraded final performance by maintaining high variability. Thus, exploration via noise introduces a limiting trade-off.

**Figure 7.**
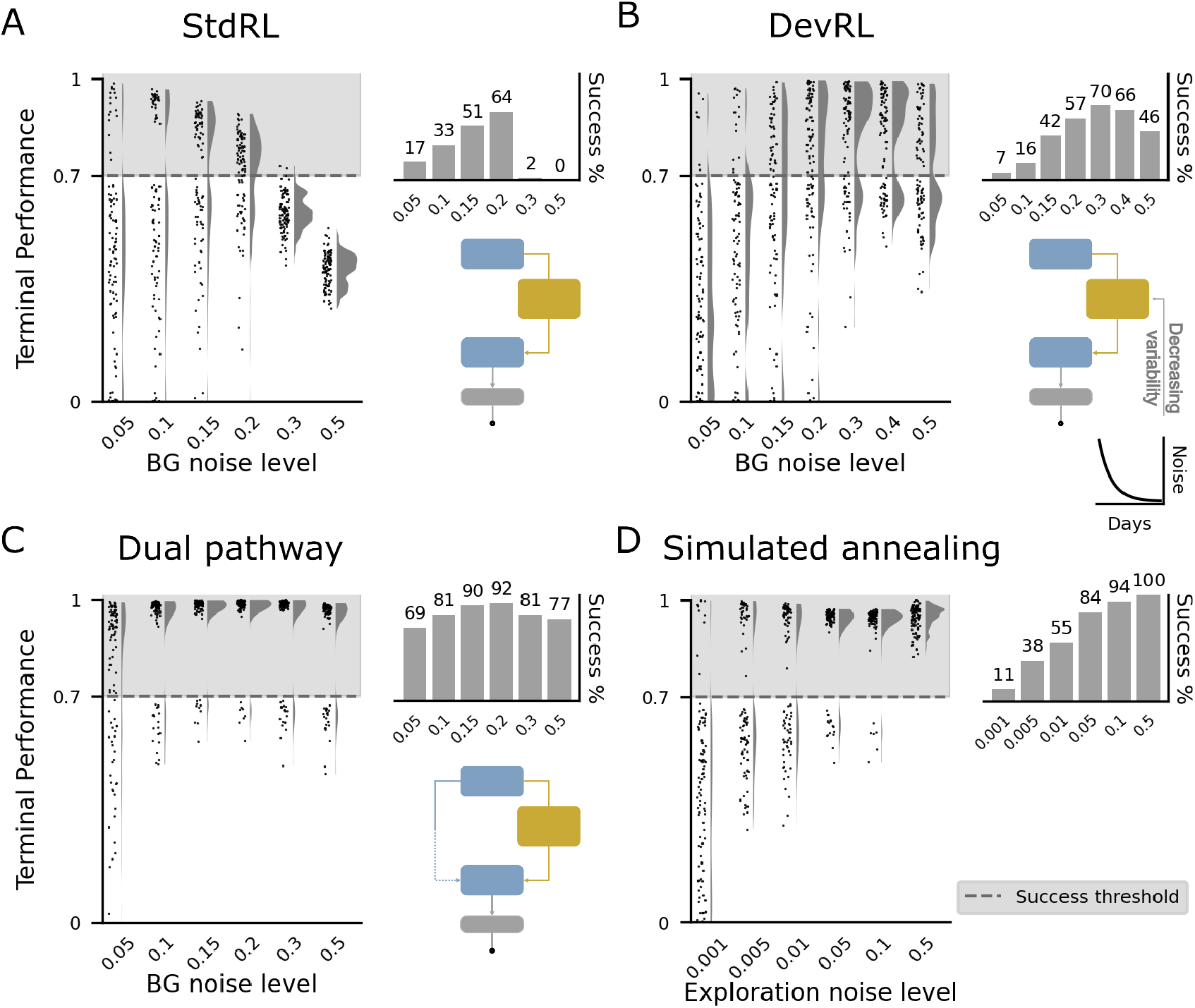
Comparison of different strategies for sensorimotor learning. In A-D, Terminal performance over 100 simulations (black dots) on artifical landscapes is depicted on the left for various levels of exploratory noise (violin plot: distribution of the terminal performance). Success rate, i.e. the proportion of simulations that learned the optimal performance (achieved terminal performance > .7), is depicted on the right for the same levels of exploratory noise. **A**. Performance of a single pathway architecture employing gradient descent based reinforcement learning with no overnight synaptic volatility (StdRL). **B**. Performance of a single pathway architecture employing gradient descent based reinforcement learning with exploratory variability reducing exponentially over development (DevRL). **C**. Performance of the dual pathway model, with overnight synaptic volatility and no explicit reduction in BG variability. **D**. Performance of a simulated annealing algorithm on sensorimotor learning on gaussian-based landscapes.

To mitigate this, we tested a variant with developmentally decreasing noise (DevRL, see Section Methods). While DevRL improved performance at moderate noise levels by enabling early exploration and later stabilization, success rates remained capped (≈ 70%, Figure 7b). Moreover, the range of noise level that allows good performance (0.3-0.4) is relatively tight and requires finetuning to be successful. At higher noise levels, performance again deteriorated, indicating that this approach does not fully resolve the exploration–exploitation trade-off.

In contrast, the dual-pathway model consistently outperformed both StdRL and DevRL across noise conditions (Figure 7c). Its combination of BG-driven variability, synaptic volatility, and gradual cortical consolidation supports broad early exploration while ensuring stable performance after convergence, without parameter fine-tuning. Finally, we compared the model to simulated annealing (see Section Methods), a standard optimization strategy that explicitly controls exploration via a temperature parameter (***Kirkpatrick et al., 1983***). Simulated annealing achieved high success rates at sufficient noise levels while maintaining good final performance (Figure 7d), owing to the temperature parameter that progressively suppresses exploratory variability over time. Despite mechanistic differences, it produces dynamics similar to the dual-pathway model.

Overall, these results show that effective sensorimotor learning requires adaptive regulation of exploration and exploitation. The dual-pathway architecture achieves this intrinsically through its biological design, providing a robust alternative to single-pathway learning strategies.

### Robustness

We finally evaluated the robustness of the dual-pathway model to parameter variations and task conditions. Increasing intrinsic BG noise initially improved performance by enhancing exploration, but excessive noise impaired convergence (Figure 8a). Notably, high success rates (>70%) were maintained across a broad intermediate range, indicating tolerance to variability in exploratory drive. Similarly, the model remained stable under substantial noise in RA, with performance degrading only at very high levels (>0.5) where learning became unstable (Figure 8b). This demonstrates robustness to motor variability.

**Figure 8.**
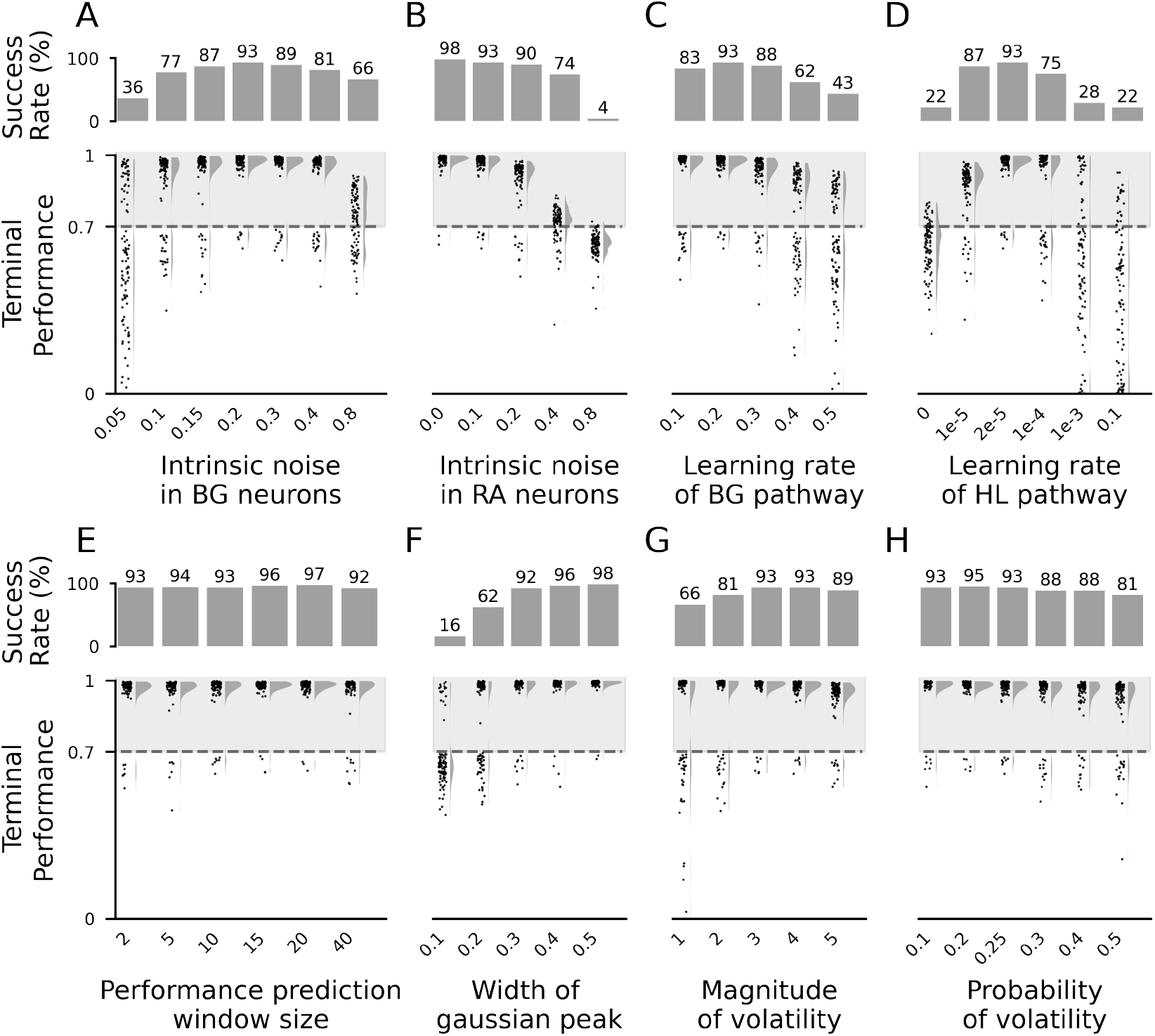
The dual pathway model is robust to fluctuations in key parameters that affect learning. Here, the performance of the model is evaluated over 100 simulations for different values of each parameter considered. **A**. Synpatic noise into BG neurons. **B**. synaptic noise into RA neurons. **C**. Learning rate for plasticity at HVC-BG synapses. **D**. Learning rate of plasticity at HVC-RA synapses. **E**. Number of renditions over which prediction of performance (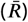) is computed (temporal integration window). **F**. Width of the global optimum gaussian in the gaussian-based performance landscapes. **G**. Scaling factor for overnight synaptic volatility at HVC-BG synapses. **H**. Probability of occurrence of overnight synaptic volatility at HVC-BG synapses.

Varying learning rates revealed complementary constraints. BG learning was robust across a range of values but became unstable at high rates (Figure 8c). Cortical learning tolerated a wide range (1*e*^−5^ to 2*e*^−4^), but very slow learning (<= 1*e*^−5^) effectively removed the cortical pathway, while overly fast learning (>= 1*e*^−3^) led to premature consolidation and suboptimal convergence (Figure 8d). Thus, efficient learning requires cortical consolidation to operate on a slower timescale than BG-driven exploration.

We also tested robustness to task structure (Figure 8e). The model reliably converged on broad optima and performed well on moderately sharp peaks, though performance declined for extremely narrow optima (62-92% success rate at gaussian peak widths of 0.2-0.3; 16% success rate at gaussian peak widths of 0.1). Learning was largely insensitive to the temporal window used to compute performance prediction error (Figure 8f), with strong performance even for minimal integration (two renditions). Finally, we examined parameters governing overnight synaptic volatility. Low volatility reduced performance, resembling purely local exploration, whereas moderate volatility improved success by enabling escape from local optima (Figure 8g). The model was relatively insensitive to volatility probability, although excessively frequent perturbations disrupted convergence by inducing persistent instability (Figure 8h).

Overall, the dual-pathway framework demonstrates robust performance across a wide range of neural, learning, and task parameters, while highlighting key constraints on the balance between exploration, consolidation, and stability.

## Discussion

We developed a computational model of sensorimotor learning grounded in the anatomy, physiology, and developmental trajectory of the zebra finch song system and singing behavior. The model integrates three key features: reinforcement learning in a cortico-BG pathway, Hebbian consolidation in a parallel cortical motor pathway, and overnight synaptic volatility in cortico-BG synapses. Our model reproduces the non-monotonic learning trajectory observed in juvenile zebra finches, the neuronal activity patterns in the network and the reported effect of circuit perturbations. Importantly, the proposed mechanism for sensorimotor learning enables reliable convergence to global optima in complex, multi-peaked performance landscapes, a setting in which standard RL approaches typically fail.

### Delayed maturation of cortical pathway implicitly steers the exploration-exploitation trade-off

Motor variability is essential early in learning, enabling exploration of movements that can later be reinforced if successful (***Sutton and Barto, 2018***; ***Wu et al., 2014***). As learning progresses, however, variability decreases and eventually impairs performance (***Cohen et al., 2007***; ***Darshan et al., 2014***; ***Pekny et al., 2015***; ***Dhawale et al., 2017***). In zebra finches, increasing similarity to the tutor song is accompanied by reduced vocal variability (***Tchernichovski et al., 2001***; ***Ölveczky et al., 2005***). Our model reproduces this transition through delayed maturation of the motor pathway (HVC–RA) and explains a key well-established experimental dissociation: the BG pathway is necessary for song learning but largely dispensable for crystallized song production (***Bottjer et al., 1984***; ***Scharff and Nottebohm, 1991***; ***Aronov et al., 2008***). This emerges from a gradual transfer of motor control from the BG pathway to the cortical pathway. Early in learning, weak HVC–RA connectivity allows BG-driven variability to dominate RA activity. Consistent with this, HVC–RA synapses emerge relatively late (>40 dph) and strengthen progressively through the sensorimotor period until 90 dph (***Herrmann and Bischof, 1986***; ***Bottjer et al., 1986***; ***Kittelberger and Mooney, 1999***; ***Garst-Orozco et al., 2014***; ***Sankar et al., 2022***). As cortical consolidation proceeds, motor patterns stabilize and dependence on BG input declines. In the model, strengthening cortical input increasingly saturates RA neurons through Hebbian consolidation of BG-guided vocalizations, progressively reducing the impact of BG variability. Consequently, BG lesions have stage-dependent effects: early lesions impair learning, intermediate lesions halt refinement, and late lesions minimally affect performance. This accounts for the decreasing influence of LMAN on RA activity and vocal output, alongside the increasing role of HVC (***Brainard and Doupe, 2000***; ***Ölveczky et al., 2005***; ***Aronov et al., 2008***; ***Kao and Brainard, 2006***). Notably, LMAN-RA synaptic input remains largely unchanged during learning (***Nordeen et al., 1992***; ***Garst-Orozco et al., 2014***), and LMAN activity itself remains variable (***Achiro et al., 2017***). Consistently cutting HVC-RA axons in older juveniles or adults simply reverts song into subsong (***Aronov et al., 2008***). Similar increases in cortical control during learning have been observed in other species (***Hwang et al., 2021***). More broadly, this subcortical-to-cortical transfer parallels phenomena such as motor-skill automatization and systems-level memory consolidation. In humans, early learning relies more on prefrontal and striatal circuits, whereas later stages depend increasingly on cortical representations (***Doyon and Benali, 2005***; ***Dayan and Cohen, 2011***). Related principles have also been proposed for hippocampal–cortical memory transfer (***Benchenane et al., 2011***; ***Maviel et al., 2004***) and cerebellar learning (***Mauk, 1997***; ***Bae et al., 2025***). In these systems, Hebbian plasticity in a direct motor or cortical pathway is thought to consolidate behaviors or memories initially acquired through a parallel circuit. The dual-pathway architecture provides a mechanistic account of these transitions grounded in pathway-specific plasticity rather than abstract learning stages.

### Role of overnight performance deterioration

Neuromuscular control of the syrinx poses a highly redundant, nonlinear sensorimotor learning problem, in which many motor configurations yield suboptimal vocal outputs (***Srivastava et al., 2015***) and transitions between them follow complex, nonlinear and discontinuous trajectories (***Tchernichovski et al., 2001***). In such landscapes, gradient-based RL readily becomes trapped in local optima. A central and counterintuitive result of our model is that overnight performance deterioration improves long-term learning. Without synaptic volatility, the system converges reliably but prematurely. Stochastic perturbations in BG synapses displaces vocal output, enabling renewed exploration and eventual convergence to the global optimum.

This provides a functional interpretation of the observation that post-sleep deterioration in juvenile zebra finches predicts better eventual imitation (***Derégnaucourt et al., 2005***). In the model, this effect arises not from passive decay but from active synaptic remodeling. Consistent with this idea, rapid spine turnover has been observed over day-long timescales in HVC (***Roberts et al., 2010***), as in the neocortex (***De Paola et al., 2006***; ***Mongillo et al., 2017***) and cortico-striatal synapses in mammals (***Shen and Dayan, 2025***; ***Kuo and Liu, 2019***). Moreover, motor learning is associated with deep structural dynamism of synaptic boutons over days, with diverse changes in neighboring synapses consistent with large stochastic changes in synaptic strength over days (***Shen and Dayan, 2025***). We further predict that the magnitude of overnight synaptic change scales inversely with within-day learning, linking daytime consolidation with nighttime exploration. This hypothesis could be tested by tracking synaptic dynamics across development or manipulating sleep. More broadly, these results support the view that sleep actively reorganizes neural circuits to enhance learning (***Rasch and Born, 2013***; ***Stickgold, 2005***). Non-monotonic learning trajectories, rather than reflecting instability, may instead be a hallmark of efficient global search in complex sensorimotor landscapes.

### Relationship to other computational models

Early RL models of song learning have demonstrated that dopaminergic signals can drive gradient ascent in motor space via BG circuits (***Doya and Sejnowski, 1994, 1998***) and have been implemented with realistic plasticity rules and spiking neuronal networks (***Fiete et al., 2007***; ***Farries and Fairhall, 2007***). Recent experimental evidence demonstrated that dopamine in the BG encode a performance error signal (***Gadagkar et al., 2016***; ***Kasdin et al., 2025***) that can drive song adaptation (***Xiao et al., 2018***). Concerning the consolidation of song adaptation initally driven by the BG-cortical circuit in the motor pathway (***Andalman and Fee, 2009***; ***Warren et al., 2011***), ***Teşileanu et al. (2017***) formalized its implementation through experimentally evidenced spike-timing plasticity at HVC–RA synapses (***Mehaffey and Doupe, 2015***), showing that the timescale of LMAN input to RA is optimal for such consolidation given the plasticity rule reveal experimentally. However, these models have generally assumed simple or unimodal performance landscapes and have not addressed the challenges posed by multiple local optima. Our model extends this framework by explicitly considering complex, multi-modal landscapes derived from a biophysical model of the syrinx. In this setting, standard RL approaches and noise-based heuristics often fail. By combining BG-driven exploration, cortical consolidation, and synaptic volatility, the model provides a mechanistic solution to escaping local optima, connecting circuit-level plasticity to the non-monotonic behavioral trajectory first described by (***Derégnaucourt et al., 2005***; ***Kollmorgen et al., 2020***) and left largely unexplained by prior models.

Our work also situates song learning within the broader landscape of sensorimotor learning theory. During sensorimotor learning, both cerebellar and BG circuits contribute to learning, albeit with learning mechanisms (***Doya, 2000***): supervised learning relying on internal models for the cerebellum (***Wolpert and Kawato, 1998***; ***Ito, 2008***), RL for the BG (***Houk and Wise, 1995***; ***Fee and Goldberg, 2011***). While cerebellar circuits may play a role in the temporal patterning of song (***Pidoux et al., 2018***; ***Ursu et al., 2026***), our model focusses on BG-related learning. Model-free RL approaches applied to continuous motor control (***Peters and Schaal, 2008***) typically assume smooth, convex performance landscapes and do not readily address the acquisition of motor programs from scratch in multi-modal landscapes typical of highly redundant motor systems with high degrees of freedom (***Bernstein, 1967***) such as the zebra finch singing apparatus (***Düring et al., 2013***). Moreover, RL implementation in neural circuits face precisely the exploration–exploitation limitations that motivated our study (***Ishii et al., 2002***; ***Humphries, 2012***).

Our model also shares conceptual similarities with optimization strategies such as simulated annealing (***Kirkpatrick et al., 1983***). Early high variability and large across-day shifts resemble high-temperature exploration, while gradual stabilization parallels cooling. However, unlike simulated annealing, which requires explicit scheduling and global memory of explored states, the dual-pathway model achieves similar dynamics through distributed, biologically plausible mechanisms. The analogy extends to mini-batch gradient descent (***Lecun et al., 1998***; ***Krizhevsky et al., 2017***), a variant of stochastic gradient descent (***Robbins and Monro, 1951***) where learning proceeds through successive subsets of data. There exists actually many different strategies (mini-batch sampler) that dictates how to build each mini-batch (***Zhao and Zhang, 2014***; ***Csiba and Richtárik, 2016***; ***Newling and Fleuret, 2016***), and reducing the batch size may improve learning (***Obando-Ceron et al., 2023***). In our model, each “day” effectively constitutes a batch, with overnight variability reshaping the sampling distribution. This may contribute to efficient convergence, although the biological implementation differs substantially from engineered algorithms. Overall, the results suggest that structural organization, i.e. parallel pathways with distinct plasticity rules and timescales, can provide a powerful and robust solution to exploration–exploitation challenges in both biological and artificial systems. Other optimization techniques used in machine learning such as simulated annealing may succeed in pathological landscapes that pose difficulties to the model proposed here. However, they are less biologically realistic as they rely on a long-term maintenance of the memory of all explored options.

### Limitations

The model incorporates several simplifying assumptions that should be addressed in future work. HVC activity is treated as fixed, whereas experimental data indicate substantial developmental changes (***Mackevicius et al., 2023***), including synaptic remodeling and neurogenesis (***Mooney, 2009***). Incorporating plastic sequence generation could reveal interactions between sensory and motor learning. Many computational models of HVC sequence generation have been proposed (including but not limited to (***Jin et al., 2007***; ***Armstrong and Abarbanel, 2016***; ***Cannon et al., 2015***)) along with mechanisms for the acquisition of the sequence pattern during sensory learning (***Fiete et al., 2010***; ***Shao et al., 2026***). The model also assumes a pre-existing dopaminergic performance signal without addressing how this evaluative signal develops. Understanding the emergence and tuning of this “critic” remains an open question.

Motor output is represented in a highly simplified, low-dimensional space. While sufficient to capture key principles, this abstraction does not reflect the full complexity of the syrinx and res-piratory system (***Düring et al., 2013***). More detailed biomechanical models could provide a richer test of the framework (***Mindlin, 2017***; ***Amador et al., 2025***). Given the difficulty of building a realistic vocal production organ, we chose to investigate the capacity of our model for a simple model of the syrinx or for various arbitrary performance landscape. Temporal structure is also simplified: each time step within a motif is treated independently, whereas real song learning involves continuous sequences with temporal dependencies. Extending the model to capture sequence-level learning and timing control is an important direction. Finally, the use of rate-based neurons omits spike-timing dynamics and detailed synaptic mechanisms. While appropriate for capturing system-level behavior, future implementations could incorporate spiking models (***Farries and Fairhall, 2007***) to better align with physiological data.

### Predictions and conclusion

Beyond reproducing known phenomena, the model generates several testable predictions. It suggests that synaptic volatility in the BG pathway—potentially mediated by spine turnover—drives overnight changes in behavior. Rapid spine formation and elimination over day-long timescales have been observed in rodent neocortex (***Holtmaat and Svoboda, 2009***) and in HVC of zebra finches (***Mooney, 2009***), although comparable measurements in Area X are currently lacking. Direct measurements of synaptic dynamics of HVC-X synapses would be required to test this hypothesis. The model also predicts that BG lesions should abolish overnight performance deterioration, and that neural activity patterns in LMAN and RA should show larger changes across nights than within days.

This work bridges behavioral, anatomical and neurophysiological evidence from songbirds and computational modeling to elucidate the mechanisms underlying sensorimotor learning. Our dual-pathway model not only aligns with existing data but also outperforms traditional RL approaches, offering a robust and biologically plausible alternative for real-world sensorimotor control. It achieves functionality analogous to optimization strategies such as simulated annealing with an implicit regulation of the exploration–exploitation trade-off through delayed maturation of the consolidation pathway. Future work could explore the applicability of these principles to other learning systems, both biological and artificial, and further refine our understanding of how the brain optimizes behavior in dynamic environments.

## Methods

Model simulations were conducted in Python. Code, parameters, and analysis scripts are publicly available at [Github](https://github.com/AtharvSuryawanshi/Dual_Pathway_RL_songbird).

### Model Architecture

Inspired by the vocal learning circuitry of songbirds (the song system) (Figure1b), we designed the model as a multilayered architecture comprising two major parallel pathways, the cortical motor pathway and the subcortical BG pathway (Figure 1c). The model consists of four layers corresponding to HVC, RA, the BG (representing the AFP), and the MC (an abstraction of the downstream motor control structures).

The first layer, HVC, serves as an input layer that provides timing and contextual information, indicating which target syllable is to be produced. The second layer, RA, generates a motor command based on input from HVC and modulatory input from the BG pathway. The third layer, BG, forms a parallel pathway between HVC and RA and receives evaluative feedback signaling performance quality. The fourth layer, MC, mimics the input to an avian syrinx and transforms RA motor commands into a low-dimensional control signal that is subsequently transformed into a syllable vocalisation using a biophysical model of the avian syrinx (***Titze and Martin, 1998***; ***Mindlin, 2017***).

In the zebra finch, the RA contains approximately 13,000–15,000 neurons per hemisphere during the early sensorimotor phase (***Herrmann and Arnold, 1991***). During this period, approximately 5,000–10,000 neurons from LMAN project to RA (***Bottjer et al., 1986***). In addition, the HVC contains roughly 23,000–40,000 neurons per hemisphere during the sensorimotor period, although only a fraction of these neurons project to RA and are active during song production (***Hahnloser et al., 2002***). These anatomical proportions and connectivity constraints informed the design of the dual-pathway model architecture.

In the model, the HVC and RA layers each comprise 100 rate-coded neuronal population units, while the BG layer comprises 50 rate-coded units. The relative sizes of these layers were chosen to approximately reflect the proportions of neurons in the corresponding nuclei of the song system, preserving a 1:2:2 ratio between the number of units in LMAN (BG layer), HVC, and RA, respectively. Both BG and RA units are modeled using sigmoidal activation functions that bound firing rates between -1 and 1 (Eqs 4, 5, 6, 7). Each unit represents the cumulative activity of a local population of excitatory and inhibitory neurons, such that its output corresponds to high (≈400 Hz for HVC, RA and ≈200Hz for LMAN) or low (≈0 Hz) effective firing rates.

The BG layer receives input from HVC (Eq 1). The RA layer receives two distinct sets of inputs (Figure 1c): a direct projection from HVC via the cortical motor pathway (Eq 2) and a modulatory projection from the BG via the subcortical pathway (Eq 3). RA activity is then transmitted to a downstream motor controller layer (MC), which transforms the RA neural command into a low-dimensional motor control signal (Eq 8).

The HVC and RA layers are fully connected, as are the HVC and BG layers. Synaptic weights in these pathways are plastic, reflecting activity-dependent plasticity at RA and BG synapses. HVC–RA weights are initialized to zero, whereas HVC–BG weights are initialized randomly from a uniform distribution over [-1, 1]. Positive weights produce a net excitatory effect on the target population, while negative weights produce a net inhibitory effect. In contrast, the BG–RA and RA–MC pathways are fully connected with fixed positive weights.

The two-dimensional motor output of the MC layer specifies the model’s sampling position in the sensorimotor space, which is subsequently transformed into a syllable-level vocalization, as described in subsection Performance Landscapes.

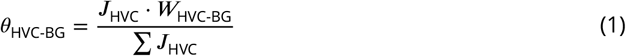

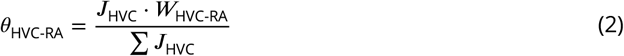

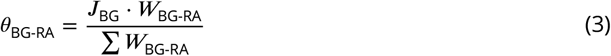

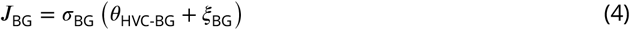

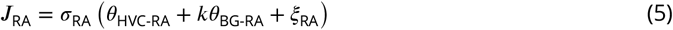

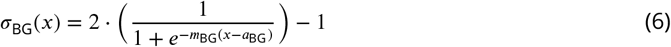

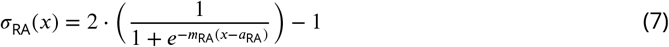

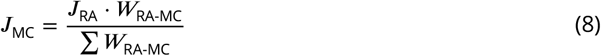

with

***J***_*x*_ - activity of the layer *x*,

***W***_*x*−*y*_ - weights connecting layer x to layer y,

*ξ*_*x*_ - intrinsic noise in layer *x*, drawn from a normal distribution

*θ*_*x*−*y*_ - input from layer x to layer y

*η* - learning rate

*k* - fixed scaling factor.

### Learning

Plastic synaptic weights in the BG pathway (***W***_HVC-BG_) are updated using a reinforcement learning rule based on covariance learning (Eq 9). Exploratory variability is introduced locally within the BG layer as intrinsic noise (*ξ*_BG_; Eq 4). In addition, the BG layer receives a scalar evaluative signal reflecting relative performance quality, quantified as the performance prediction error (PPE). The PPE is defined as the difference between the performance evaluation on a given trial (*R*) and the expected performance evaluation (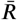; Eq 10). In contrast, plastic synaptic weights in the cortical motor pathway (***W***_HVC-RA_) are updated using a Hebbian learning rule, whereby coincident activity in pre- and post-synaptic units leads to potentiation of the corresponding synapse (Eq 12). All plastic synaptic weights are bounded within the range [-1,1].

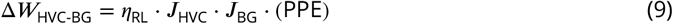

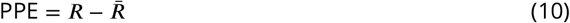

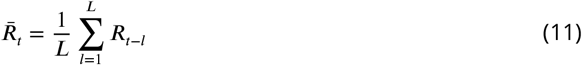

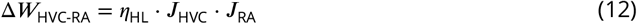

Further, the reinforcement learning driven by the BG pathway is supplemented with behavioral features drawn from empirical evidence. Specifically, a daily deterioration in song performance has been shown to occur post-sleep in juvenile birds during the sensorimotor period. This post-sleep deterioration is implemented as a fluctuation of the HVC-BG weights at the end of a day (Eq 13), which results in a displacement of the BG output on the following day. The magnitude of the overnight volatility in the HVC-BG synapses is determined stochastically (*ξ*_night_) from a uniform distribution [*−*1, 1] and is scaled inversely with the change experienced by the HVC-BG weights over the previous day (*p*) (Eq 14).

Note: For comparisons with the model when lacking overnight jumps (Fig 5), this overnight synaptic volatility was omitted, while all other aspects of the model and learning rules were kept identical.

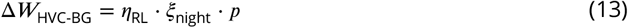

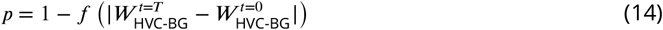

where *f* is a sigmoid function with an optimum threshold and slope.

### Simulation

Within each simulation, the model learns a song motif composed of four syllables in the task with the syrinx-based performance landscapes, and 20 sub-syllabic notes (4 syllables × 5 sub-syllabic notes) of 10ms each in the task with the artificial performance landscapes, over a 60-day sensorimotor period (Figure 1a). Each day, the model performs 1000 renditions of the song motif, with each rendition consisting of the syllables or sub-syllabic notes generated in a fixed temporal sequence. For each rendition of a syllable or sub-syllabic note, a performance metric, indicating its similarity to the corresponding target syllable or note, is obtained from the performance landscape corresponding to the target syllable or note, and used to compute the performance prediction error (PPE).

**Table 1.** Model parameters and their values used in all simulations unless otherwise stated.

| Parameter |  | Value |
| --- | --- | --- |
| Reinforcement Learning rate | $\eta_{RL}$ | 0.2 |
| Hebbian Learning Rate | $\eta_{HL}$ | $2 \times 10^{-5}$ |
| BG noise | $\xi_{BG}$ | 0.2 |
| RA noise | $\xi_{RA}$ | 0.1 |
| Performance prediction window size | $L$ | 10 |
| Landscape density |  | Moderate |
| Global optimum gaussian width |  | 0.3 |
| Magnitude of volatility |  | 0.4 |
| Probability of volatility |  | 0.25 |

### Performance Landscapes

We evaluated the model using two types of performance landscapes: syrinx-based and Gaussian-based landscapes (Figure S1). For the syrinx-based landscapes, the MC layer generates a motor command that is mapped to a vocal output, which is then evaluated relative to a target tutor song. This transformation and comparison are implemented using a biophysical model of the avian syrinx, allowing motor commands to be translated into vocalizations whose similarity to the tutor template defines performance quality. The two-dimensional scalar output of the MC layer, ***J***_MC_, is transformed into input signals for the syrinx model according to Eq 15 and Eq 16. Specifically, the MC layer modulates two control signals, 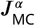 and 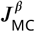, corresponding to air-sac pressure, *α*(*t*), and syringeal labial tension, *β*(*t*), respectively. These pressure and tension waveforms drive oscillations in the trachea, generating a syllable vocalization as a pressure waveform of duration ***T*** = 50 ms (Supplementary figure S1a; ***Amador and Margoliash (2013***)). A spectrogram is constructed from this oscillatory pressure waveform and compared to a target syllable selected from the vocal repertoire of zebra finches. Vocal similarity is quantified using a similarity metric (*s*) based on the correlation coefficient between the normalized spectrograms of the generated vocalization (***S***_*X*_) and the target template (***S***_***T***_) (Eq 17).

To construct the performance landscape, we simulated vocalizations over a range of MC outputs— corresponding to different combinations of pressure and tension parameters—and computed the similarity metric for each vocalization relative to the target syllable. The resulting performance landscape is represented as a two-dimensional matrix that captures vocal similarity across the pressure–tension input space for a given target syllable. Using four distinct target syllables, we generated four corresponding syrinx-based performance landscapes. As shown in supplementary Figure S1b, the resulting performance landscapes exhibit multiple global optima, reflecting the inherent redundancy of the syrinx, whereby distinct combinations of pressure and tension can produce acoustically similar vocal outputs. In addition to these global optima, the landscape contains numerous local optima distributed across the sensorimotor space, arising from the nonlinear biomechanics of sound production.

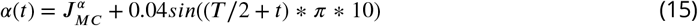

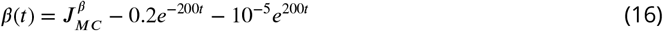

where the output of the MC, ***J***_*MC*_ *∝* [***J*** ^*α*^, ***J*** ^*β*^], ***T*** : duration of the vocalisation and *t* ∈ [0, ***T***].

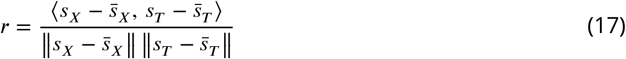

Here, *s*_*k*_ denotes one-dimensional vector obtained by flattening the two-dimensional matrix ***S***_*k*_, 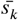 denotes its mean and *r* corresponds to their Pearson correlation coefficient.

Second, to generate performance landscapes with systematically varying complexity, we adopted a direct computational approach. We constructed continuous performance landscapes by mapping the two-dimensional motor space (***J***_MC_) onto a one-dimensional scalar performance evaluation. Each performance landscape was generated as the maximum over a set of two-dimensional Gaussian distributions, each centered at a distinct randomly sampled location in motor space and characterized by a specific width. Similar to the syrinx-based performance landscapes, the superposition of these distributions yields landscapes with multiple local optima alongside a single global optimum. Landscape complexity was controlled by varying the number, spatial arrangement, and widths of the Gaussian components. Local optima were constrained to reach a maximum performance value of 70% of that at the global optimum, and all landscapes were normalized such that the global optimum yielded a performance metric of 1. Based on the density of local optima, landscapes were grouped into four classes of : low (2-5), moderate (3-9), compact (8-18) and high (11-25) number of local optima. All simulations were run on landscapes the moderate density of local optima, unless otherwise stated. Figure S1d shows representative examples from each class. These Gaussian-based landscapes allow us to systematically assess the versatility and robustness of the model across a wide range of task complexities.

### Performance Metrics

Model performance was quantified using two metrics: terminal performance and success rate. The terminal performance of a syllable was defined as the mean performance evaluation obtained from its corresponding performance landscape over the last 100 renditions on the final day of learning. This metric captures the imitation quality of the learned motor output after learning has stabilized.

For the Gaussian-based performance landscapes, the global optimum was assigned a performance value of 1, while all local optima were constrained to reach a maximum performance value of 0.7. Accordingly, a simulation was considered successful if it achieved a terminal performance exceeding 0.7. For the syrinx-based performance landscapes, we observed that the highest peak outside the global optimum had an associated performance value of approximately 0.52, whereas the global optima reached a value of 1. To maintain consistency across landscape types, we apply the same success threshold of 0.7 for syrinx-based simulations. The success rate for a given experimental condition was defined as the proportion of simulations that exceeded the success threshold relative to the total number of simulations.

Motor variability is calculated as standard deviation of the output on each dimension over the day. End-of-day metrics are calculated as the mean of the metric over the last 100 renditions of the day. Start-of-day metrics are calculated as the mean of the metric over the first 100 renditions of the day.

### Experiments

To isolate the contribution of the basal ganglia (BG) pathway to sensorimotor learning, we implemented a lesion-like manipulation by silencing BG output to RA. Specifically, the projection from the BG pathway to the RA layer was inactivated, effectively removing LMAN-mediated input to RA while leaving the cortical (HVC–RA) pathway intact. This manipulation allowed us to assess model performance when motor production and learning were governed solely by the cortical pathway. Lesions were applied at different stages of sensorimotor learning to examine developmental effects. We performed the manipulation at three time points corresponding to before learning begins (40 dph), during learning (70 dph), and post learning (100 dph). Following the lesion, simulations were continued without further modification to learning rules or parameters, and performance metrics were evaluated as described above. We performed an analogous manipulation targeting the cortical pathway, inactivating the HVC–RA projection while leaving the BG pathway intact, to assess learning when driven solely by BG input.

### Illustrations

To illustrate spiking activity corresponding to the scalar firing rates generated by the model, we create raster plots in Figure 1, 4. Spike trains were simulated where the firing rate (r) generated by the model for every 10 ms time bin is linearly mapped onto physiologically observed rate of firing of the particular neuron (R = 400 Hz for HVC, RA neurons and R = 200 Hz for LMAN neurons). For each time bin of Δ = 1*ms* during song, probability of a spike being generated was determined as *p ∝ r*.***R***.Δ*t*. A refractory period of 1 ms was enforced after every occurrence of a spike.

### Benchmark Algorithms

To compare the performance of the dual-pathway model with established computational approaches, we constructed a single-pathway framework implementing different reinforcement learning (RL) strategies. In this framework, a single pathway injects exploratory noise, receives performance evaluation, updates synaptic weights, and directly governs motor output. Operationally, this corresponds to lesioning the cortical motor pathway and allowing the BG-like pathway to exert sole control over motor output.

First, we evaluated the single-pathway framework using a standard reinforcement learning approach (StdRL). In this condition, learning is governed by the gradient-descent-based update rule applied in the BG pathway of the dual-pathway model (Eq 9).

Second, we tested a modified reinforcement learning strategy with developmentally decreasing noise (DevRL). In this approach, the magnitude of exploratory noise injected into the single pathway is reduced exponentially over the course of learning (Eq 19) such that the final noise is around 10% of the initial noise.

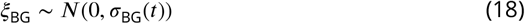

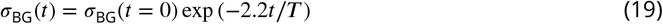

Third, we compared the dual-pathway model with simulated annealing (SA), a probabilistic optimization technique commonly used to identify global optima in complex search spaces. In this framework, transitions to lower-reward states are permitted with a probability determined by an acceptance function M, which is determined by an exponentially decreasing temperature parameter, scaled by the difference in performance evaluation between successive iterations (Eq 20, 21, 22). This temperature parameter explicitly controls the balance between exploration and exploitation over learning.

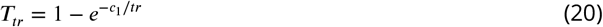

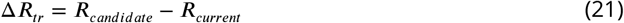

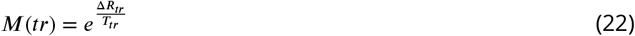

where Δ***R***_*tr*_ denotes the difference in performance evaluation between a randomly chosen candidate motor output, ***R***_*candidate*_ and the current motor output ***R***_*current*_. The candidate motor output is sampled within a range corresponding to *η*_*rl*_ * *ξ* (Eq 9), ensuring a step size comparable to that used in the preceding reinforcement learning algorithms.

## Acknowledgments

We would like to thank Alexis Dubreuil for thoughtful discussions and Manfred Gahr from the Max Planck Institute for Biological Intelligence for technical support and funding. This work has been supported by funding from the French Agence Nationale de la Recherche (grant number ANR-17-CE24-0036) to NR and (20-CE37-0023-02, 22-NEUC-0002-01) to AL, as well as Fonds pour la Recherche Médicale (EQU202203014688) to AL.

## Supplementary

**Figure S1.**
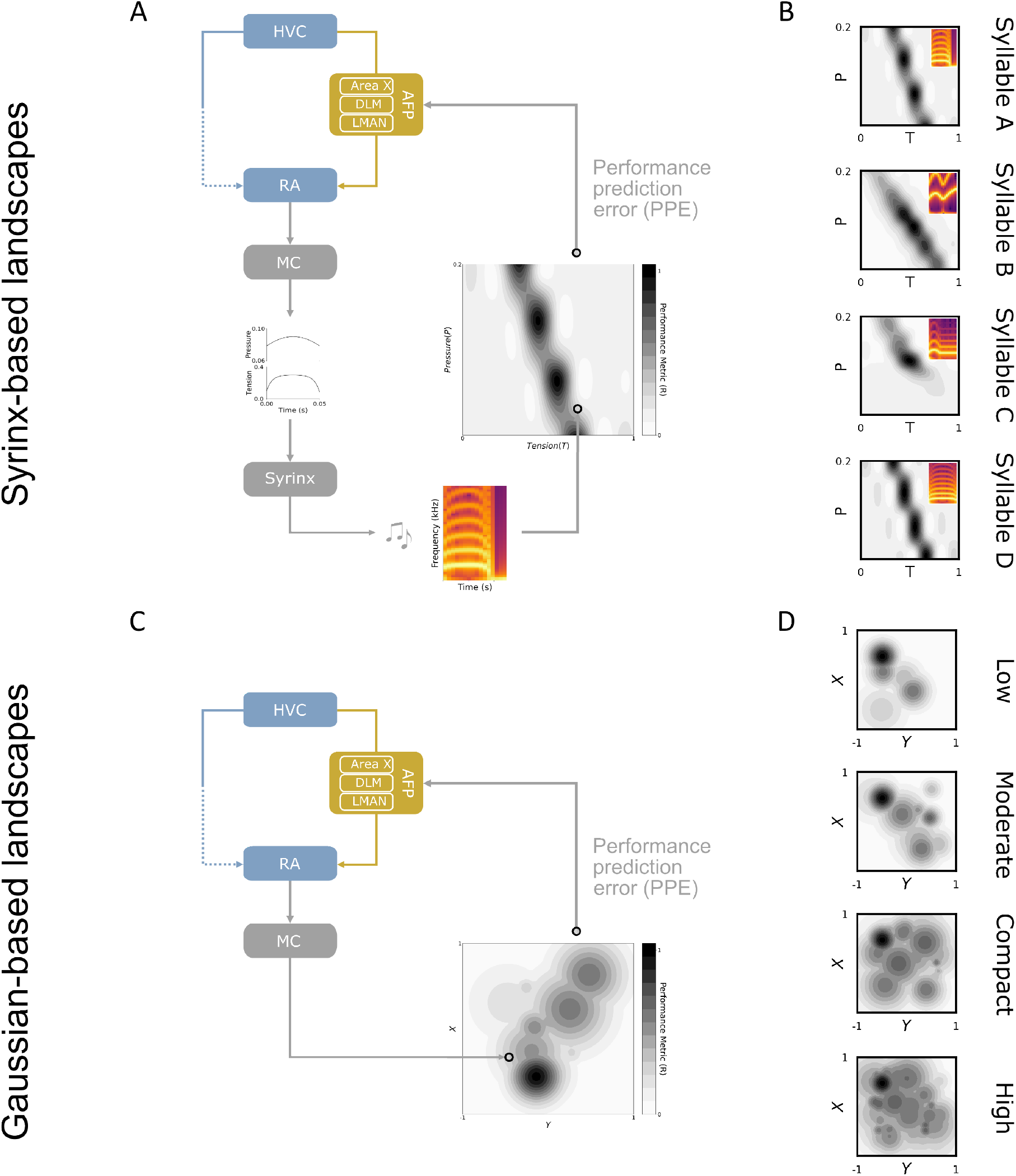
Construction of sensorimotor performance landscapes. **A**. Interface of model with syrinx-based landscapes. The neuronal activity pattern in RA is reduced in dimension at the MC layer. The output from the MC layer modulates 50 ms tension and pressure waves, which is fed as input to a biophysical model of the syrinx to produces zebra finch-like vocalizations. For each output of the model, the produced vocalization is compared with a target syllable to generate a performance metric (grey heatmap). Thus, the performance landscape represents the imitation quality for every model output. **B**. Four sample performance landscapes generated using four target zebra finch syllables (inset). The landscapes have a few global optima (performance metric ≈ 1) and several local optima (performance metric < .7). **C**. Interface of model with gaussian-based landscapes. The output from the MC layer is directly used as coordinates on these sensorimotor performance landscapes. **D**. Examples of gaussian-based landscapes from each density category. The landscape is generated by superimposing several gaussian hills. The superimposition results in several local (performance metric < .7) and one globally optimal solution (performance metric = 1).

## Footnotes

1 All summary statistics are reported as median [interquartile range (IQR)], unless otherwise stated.

